# Ecological interactions in breast cancer: Cell facilitation promotes growth and survival under drug pressure

**DOI:** 10.1101/2021.02.01.429214

**Authors:** Rena Emond, Jason I. Griffiths, Vince Kornél Grolmusz, Aritro Nath, Jinfeng Chen, Eric F. Medina, Rachel S. Sousa, Frederick R. Adler, Andrea H. Bild

## Abstract

The interplay of positive and negative interactions between drug sensitive and resistant cells influences the effectiveness of treatment in heterogeneous cancer cell populations. In a study of isogenic estrogen receptor positive (ER+) breast cancer cell lineages sensitive and resistant to ribociclib-induced CDK4/6 inhibition in mono- and co-culture, we find that sensitive cells grow and compete more effectively in the absence of treatment. During treatment with ribociclib, sensitive cells survive and proliferate better when grown together with resistant cells than when grown in monoculture, termed facilitation in ecology. Both liquid chromatography-tandem mass spectrometry (LC-MS/MS) assays and single cell RNA-sequencing (scRNAseq) indicate that resistant cell production of estradiol, a highly active estrogen metabolite, is the mechanism of facilitation. Higher estradiol production by resistant cells drives a shift in sensitive cell phenotype to a resistant cell state in coculture. Adding estradiol in monoculture provides sensitive cells with increased resistance to therapy, and cancels facilitation in coculture. Mathematical modeling quantifies the strength of competition and facilitation and predicts that blocking facilitation has the potential to control both resistant and sensitive cell populations and inhibit the emergence of a refractory population.

## Introduction

Breast cancer is the most common cancer worldwide and the second leading cause of cancer death in American women. The majority (∼80%) of these breast tumors are estrogen receptor positive (ER+), and the majority of metastatic patients who die from their cancer have this breast cancer subtype (1-3). In these tumors, estrogen receptor activity leads to cancer cell proliferation through CDK4/6 activation and cell cycle progression (4-6). In order to target both upstream ER and downstream CDK4/6 signaling for cancer control, the combination of CDK inhibitors with endocrine therapy has been used successfully in metastatic ER+ breast cancer, and to a moderate extent in earlier-stage, non-metastatic breast cancer (7-12). However, tumors can develop resistance to both single and combination endocrine and cell cycle therapy regimens (7, 8, 10-15). Understanding the underlying causes of resistance to endocrine and cell cycle therapies is a critical area of research for this major cancer subtype and cause of death in women.

Cancerous tumors consist of genetically and phenotypically heterogeneous cells (16). Despite advancements in understanding tumor heterogeneity, this feature is not used in cancer treatment strategies. ER+ breast cancer is often polyclonal and phenotypically heterogeneous, with co-existing cells of different levels of estrogen receptor expression and different levels of estrogen addiction (17). Heterogenous tumors create three major obstacles for treatment. First, cells can have different susceptibilities to treatment, meaning that even a targeted treatment with high efficacy can fail to kill or inhibit a subset of cancer cells (18, 19). Second, this differential survival and proliferation can promote continued evolution of tumor resistance during drug therapy (16, 20). Third, heterogeneous cell populations create the possibility for interactions among cells which can alter responses to treatment and patient outcomes (21-27). In this case, understanding how subpopulations communicate could help design therapy strategies that disrupt and/or exploit these interactions for clinical benefit.

This study explores the dynamics of cell interactions, including cooperation and competition, in ER+ breast cancer. Cooperation takes multiple forms and seldom exists in isolation from competition (28, 29). We investigate **facilitation**, the widely-used ecological term describing cooperative cases where competitors of one type benefit those of another under appropriate conditions (30). For example, facilitation emerges among plants competing for water when one plant gains from water that leaks from the roots of a deeper-rooted neighbor (31). This coincidence of competitive and facilitative interactions requires detailed mathematical models to disentangle their concurrent effects, and experiments to identify mechanisms.

To investigate the type, strength and mechanism of cell interactions in heterogeneous cancer populations during treatment, we developed ER+ breast cancer cell lineages that are sensitive or resistant to CDK4/6 cell cycle inhibition (ribociclib), a standard therapy used to treat this cancer. Using mathematical models of population growth and interaction, we found that ribociclib resistant cells facilitate sensitive cancer cell growth during drug treatment. Experimental analysis using liquid chromatography-tandem mass spectrometry (LC-MS/MS) assays and Western blot analysis showed that resistant cells secrete estradiol, which given the sensitive cells’ high dependence on estrogen for proliferation, leads to their growth promotion in coculture. Single-cell RNA-sequencing (scRNAseq) analysis revealed that during coculture sensitive cells acquire the traits of resistant cells and increase proliferation levels. Examining mono- and coculture growth trajectories across a broader range of drug doses, we parameterized a mechanistic model of facilitation. This consumer-resource model revealed that facilitation impairs cell cycle inhibition therapy during the rapid cancer population growth phase and predicts that blocking facilitation can jointly control resistant and sensitive populations.

These studies uncover a novel mechanism of resistance in which increased levels of local estradiol, the active estrogen metabolite, is produced by a subset of cancer cells, leading to cooperative survival and growth of normally therapy-sensitive cells. By understanding cooperative interactions like facilitation, we have the potential to better control coexisting populations within a heterogenous tumor and inhibit the emergence of resistant phenotypes.

## Results

### Sensitive and resistant cell growth and treatment effects

We developed an *in vitro* model to study ER+ breast cancer cell interactions under selective drug pressure (see Methods). To generate isogenic lineages, we applied ribociclib to cell cultures for 6-9 months until resistance developed compared to control drug sensitive parental lineage not exposed to treatment (32). These resistant and sensitive lineages, derived from CAMA-1 ER+ breast cancer cell lines, were labeled with lentivirus to express a fluorescent protein for monitoring each population’s growth when cocultured, and cell counts were calculated as a measure of spheroid area and fluorescence intensity integrated into fitted growth equations (**Supp. Fig. 1**). The procedure was repeated for two other cell lines, MCF7 and MCF7/LY2 (MCF7 cell line resistant to antiestrogen) (**Supp. Fig. 2**) (33, 34). When grown as 3D spheroids in monoculture, untreated sensitive CAMA-1 cell populations grew more quickly than resistant cells. In contrast, while under both 200nM and 400nM concentrations of ribociclib, resistant cells grew more quickly than sensitive cells **(Fig. 1a-b)**. CAMA-1 sensitive cell proliferation was inhibited at higher ribociclib concentrations (200nM and 400nM) whereas resistant cell proliferation was inhibited much less **(Fig. 1c-d)**. The log fold change of the sensitive cell population between days 4 and 14 is reduced by a factor of 7.8 by 400nM ribociclib treatment, and that of the resistant cell population by a factor of 1.25. Similarly, MCF7/LY2 sensitive cell proliferation decreased with an increase in ribociclib treatment compared to resistant lineages (**Supp. Fig. 2**).

**Figure 1.**
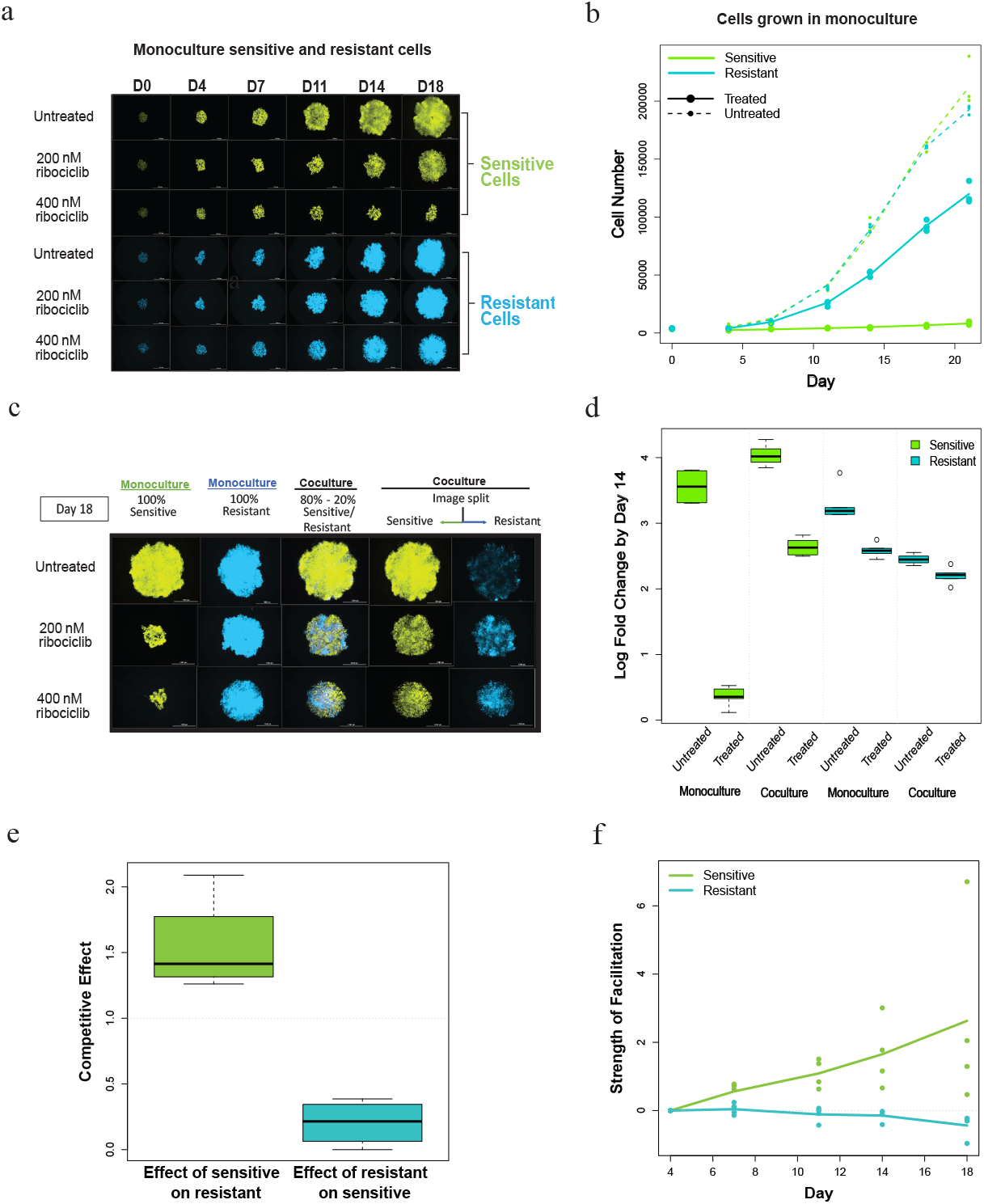
Sensitive and resistant cell growth, treatment effects, and ribociclib induced facilitation. **a**, CAMA-1 spheroids of 100% monoculture sensitive (venus, green) or resistant cells (cerulean, blue) cultured in untreated control, 200 nM ribociclib, and 400nM ribociclib treated medium for 18 days. Images taken at Day 18. **b**, Growth curves of CAMA-1 untreated and ribociclib treated (400 nM) sensitive and resistant cell populations. **c**, CAMA-1 spheroids of different cell compositions (100% sensitive - green, 50% sensitive-50% resistant, 100% resistant - blue) cultured in untreated, 200 nM ribociclib, and 400 nM ribociclib treated medium for 18 days. Images taken at Day 18 **d**, Box plot of CAMA-1 log fold change of untreated and ribociclib treated (400 nM) sensitive or resistant cells from day 0 to day 14 in monoculture and coculture (n=3). In monoculture, sensitive cells have a significantly higher reduction in growth than resistant cells (p<0.0001). Sensitive cells have a significantly lower growth reduction in coculture than in monoculture (p < 0.0001). Resistant cells show no significant effect (p=0.2) and a significantly smaller change in growth due to treatment in coculture (p=0.005). **e**, Box plots of competitive effect of sensitive (S) cells on resistant (R) cells and vice versa for CAMA-1 cells (p=0.0023 with a t test, n=3). **f**, Facilitation measured as log observed growth relative to expected growth averaged over all replicates for CAMA-1 cells (p < 0.001 for interaction of cell type with day using a linear model of the log of observed over expected cell number as a function of time).

### Ribociclib tolerance provided by facilitation of sensitive by resistant cells

To investigate whether cell interactions alter response to therapy, we compared growth of monocultured and cocultured sensitive and resistant cells treated with different doses of ribociclib over 18 days. We use Lotka-Volterra competition models (35) to quantify cell interactions and to predict the joint effect of treatment and coculture if one cell type does not alter the drug tolerance of the other. In particular, we determine the expected effect by estimating the cost of treatment and the cost of competition and predicting the combined effect by multiplying the reductions in the growth rate and the carrying capacity (**Supp. Fig. 8a** and Methods). This expected combined effect was then compared with observed growth. Observed growth greater than expected growth of sensitive cells in treated coculture indicates facilitation.

Using the competition model, we find that sensitive cells strongly suppress resistant cells in untreated cocultures, with a competitive effect 50% larger than that of resistant cells on themselves (scaled to 1.0 in these models). In contrast, resistant cells had almost no detectable competitive effect on sensitive cells (**Fig. 1e**). Facilitation was measured as the log observed growth relative to expected growth. Based on the individual effects of ribociclib and coculture, sensitive cells are expected to grow slightly more slowly in ribociclib treated coculture than in treated monoculture due to competition. Instead, we observed markedly increased growth showing facilitation of sensitive cells by resistant cells (facilitation under 200nM ribociclib = 0.973; facilitation under 400nM ribociclib = 2.39) (**Fig. 1f**). Reciprocally, resistant cells grew more slowly than expected in treated coculture due to the increased suppression by the sensitive cells they facilitate (facilitation under 200nM ribociclib = -0.055; facilitation under 400nM ribociclib = -0.227). Despite differences in competitive interactions, facilitation of sensitive cells was also observed in the additional ER+ cell lines, MCF7 (facilitation under 2.4uM ribociclib = 0.239; facilitation under 5uM = 0.4) and MCF7/LY2 (facilitation under 3uM = 0.659; facilitation under 5uM = 0.479), with MCF7/LY2 more closely resembling CAMA-1 cell growth and facilitation (**Supp. Fig. 2**).

### Mechanisms of facilitation under ribociclib treatment

To determine the mechanisms driving facilitation during ribociclib treatment, we tested whether resistant cells metabolize ribociclib more effectively than sensitive cells, reducing the ribociclib concentration and allowing increased proliferation of cocultured sensitive cells. Using an optimized HPLC/MS method, we found that the ribociclib concentration was not decreased more in media incubated with resistant spheroids than with sensitive spheroids, and that ribociclib remained at high doses (**Supp. Fig. 3**).

Alternatively, ribociclib resistant cells may release signaling molecules that enhance the growth of sensitive cells under drug treatment. To test this hypothesis, we compared proliferation of sensitive cells when supplemented with conditioned media transferred from spheroids of different compositions (100% sensitive, 50%-50% sensitive/resistant, 100% resistant) with or without treatment (0 or 400nM ribociclib). Conditioned media originating from spheroids containing resistant cells increased proliferation of sensitive cells significantly more than conditioned media produced by sensitive cell spheroids, with the largest benefit from cells that were themselves treated **(Fig. 2a)**, indicating that resistant cells secrete signaling molecules that provide pro-growth benefits for sensitive cells under drug pressure. Comparisons of sensitive cell proliferation identified no effect of exosomal transfer on sensitive cell growth under treatment (**Supp. Fig. 4**).

**Figure 2.**
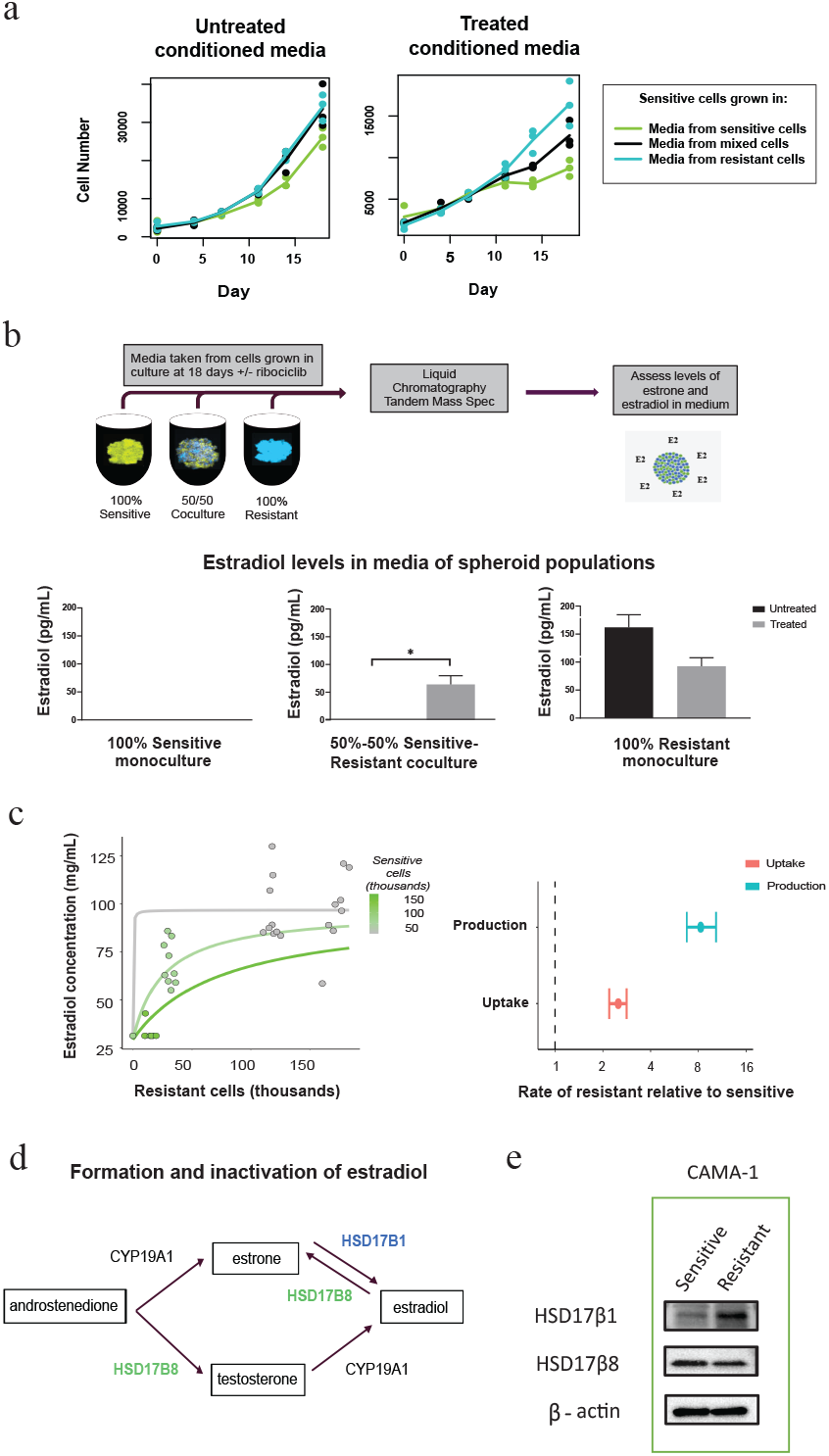
Underlying biological processes driving facilitation in response to ribociclib treatment. **a**, Growth of CAMA-1 spheroids under drug pressure with supplemented conditioned media from spheroids of different compositions. Sensitive and resistant cells were plated by the indicated composition of sensitive and resistant cells in a 3D spheroid format and underwent exposure to untreated or 400nM treated medium to produce conditioned media. 100% sensitive spheroids under drug pressure were then supplemented with different concentrations of conditioned media. Normalized cell counts of sensitive cells supplemented with untreated or treated conditioned media generated from different spheroid compositions. **b**, Liquid chromatography-tandem mass spectrometry (LC-MS/MS) assay for estradiol detection performed on CAMA-1 untreated or ribociclib 200nM treated samples of the following compositions at Day 21: 100% sensitive, 50% sensitive – 50% resistant, 100% resistant samples. **c**, Estradiol concentration by resistant and sensitive cell abundance; Estradiol production rates of CAMA-1 ribociclib sensitive and resistant cells and sensitive cell uptake of estradiol (relative to resistant cells) **d**, diagram depicting CAMA-1 resistant and sensitive cell contributions of estradiol metabolism enzymes HSD17β1 and HSD17β8. **e**, Western blot analysis of HSD17β1, HSD17β8, and β-actin (control) in monoculture, untreated sensitive and resistant CAMA-1 cells.

To identify the molecules secreted by resistant cells that may promote growth under drug pressure, we measured the concentrations of a broad range of growth-promoting factors in the media of cultured spheroids. Custom multiplex cytokine analysis for detection of known growth factors (TGFβ 1-3, TGFα, EGF, and FGF2, FGF21, FGF23) showed no significant differences in levels in cocultured sensitive and resistant cells (**Supp. Fig. 5a and 5b**). However, concentration of estradiol, a potent estrogen compared to estrone, significantly differed between resistant and sensitive spheroids. We used liquid chromatography-tandem mass spectrometry (LC-MS/MS) assay to detect estrone (**Supp. Fig. 5c**) and estradiol concentration under no treatment or 200nM ribociclib treatment **(Fig. 2b)**. Estradiol was not detectable in media from 100% sensitive cell cultures in either treatment; in contrast, samples from cultures containing resistant cells showed measurable levels of 64 pg/mL. Estradiol production and uptake by resistant and sensitive cells was estimated by fitting a Michaelis-Menten model (35) and accounting for limits of detection (**Fig 2c, left panel, Supp. Inf**.). Resistant cells are estimated to produce approximately 8 times more estradiol and use only 2.5 times more than sensitive cells **(Fig. 2c, right panel)**, generating a substantial source of the growth-promoting hormone.

Given the higher estimation of estradiol produced by resistant cells, we investigated the estrogen biosynthesis pathway and possible differences in the levels of enzymes involved in estradiol metabolism between sensitive and resistant cells. In general, breast cancer cell lines express a significantly lower level of estrogen synthetase, CYP19A1 (Aromatase), than non-cancer cell lines, with ER+ breast cancer lines expressing much lower aromatase mRNA levels than ER-lines (36, 37). However, differences in protein levels were detected in a subset of the core estradiol metabolism conversion enzymesHSD17β1 (involved in the conversion of estrone to estradiol, the more active estrogen metabolite), and HSD17β8 (an oxidative enzyme responsible for inactivating estradiol) (**Fig. 2d)**. Western blot analysis of these conversion enzymes indicates higher detectable levels of HSD17β1 in CAMA-1 resistant cells and higher levels of HSD17β8 in sensitive cells (**Fig. 2d, Supp. Fig. 6**). Similarly, levels of HSD17β8 were found to be decreased in resistant MCF7/LY2 and MCF7 cells (**Supp. Fig. 6**), suggesting that increased levels of estradiol may be driven through multiple mechanisms. While MCF7/LY2, like CAMA-1, exhibited higher levels of HSD17β1 in resistant cells, a distinction was not found between sensitive and resistant MCF7 cells, consistent with the lower facilitation in MCF7 cells than in MCF7/LY2 and CAMA-1. Experiments in media deprived of androgens from charcoal stripped fetal bovine serum produced total growth inhibition in both resistant and sensitive cells (**Supp. Fig. 7**). These analyses support a model of facilitation whereby ribociclib resistant cells produce excess estradiol via an androgen synthesis pathway with increased HSD17β1, countering sensitive cells’ inactivation of estradiol via HSD17β8, and ultimately promoting sensitive cell growth during treatment (**Fig. 2e**).

Applying ribociclib treatment in coculture reveals facilitation of sensitive cells by resistant cells. Furthermore, the degree of facilitation increases in a dose dependent manner (**Fig. 3c**). Our experiments indicate that resistant cells facilitate sensitive cells by increasing the level of local estradiol. To test this mechanism, we added estrogen pathway modifiers to cells in monoculture and coculture, both with and without ribociclib treatment, focusing on estradiol itself and the ER antagonist fulvestrant, a selective estrogen receptor degrader (SERD). We measured strength of facilitation in sensitive (**Fig. 3c, 3d**) and resistant cells (**Supp. Fig 8g, 8h**). We tested two specific predictions.

**Figure 3.**
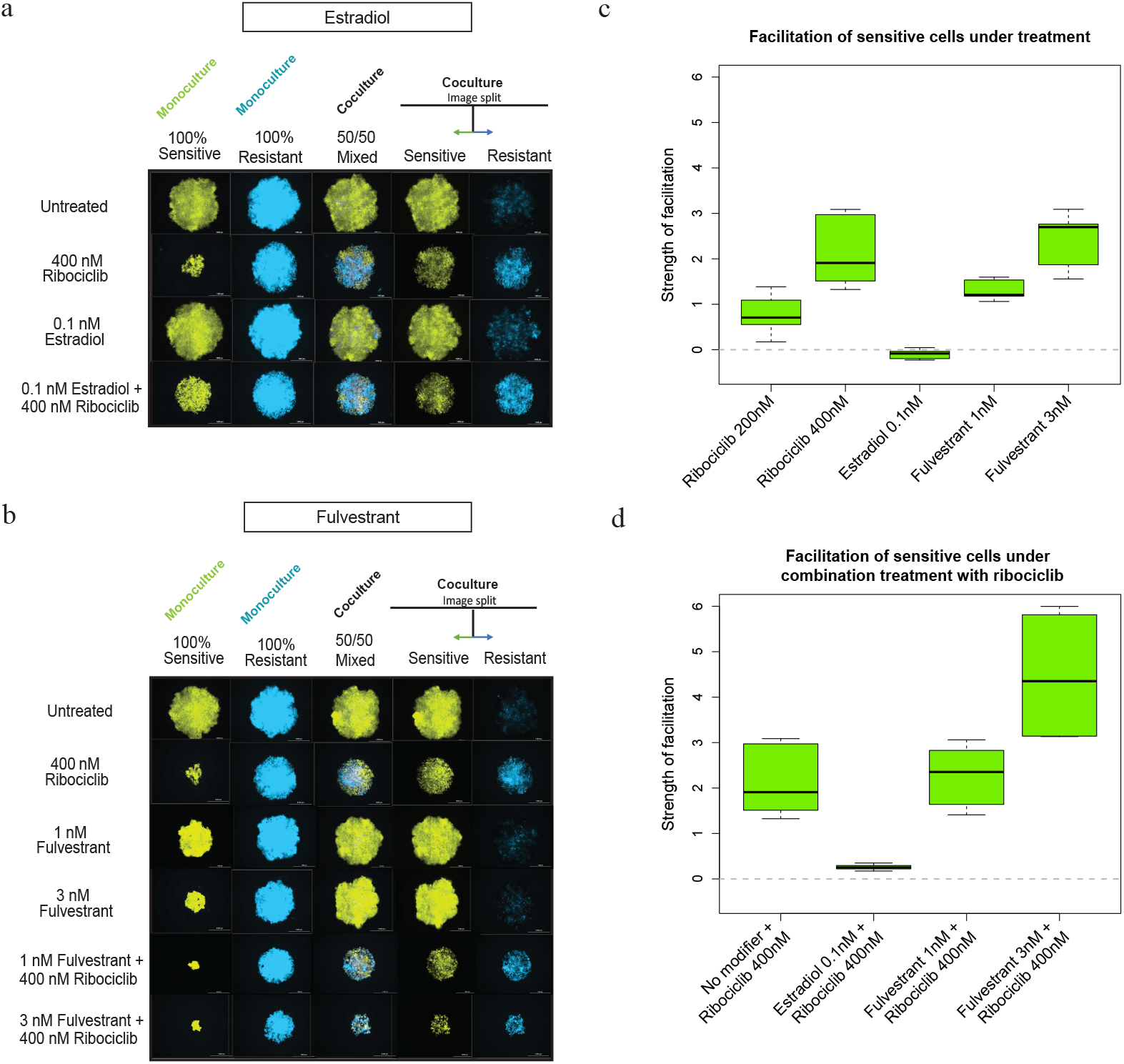
Modifying endocrine signaling pathways affects spheroid growth and facilitation. **a**, CAMA-1 spheroids of different cell compositions (100% sensitive - green, 50% sensitive- 50% resistant, 100% resistant - blue) cultured in untreated, 400nM ribociclib, 0.1nM estradiol, or combination 400nM ribociclib with 0.1nM estradiol treated medium for 18 days; images taken on day 18. **b**, CAMA-1 spheroids of different cell compositions (100% sensitive - green, 50% sensitive- 50% resistant, 100% resistant - blue) cultured in untreated, 400nM ribociclib, 1nM or 3nM fulvestrant, or combination 400nM ribociclib with 1nM or 3nM fulvestrant treated medium for 18 days; images taken on day 18. **c**, Sensitive cells with single treatment (n=3). When treated with ribociclib or fulvestrant, sensitive cells in coculture with resistant cells are significantly facilitated (p<0.0001) each with higher facilitation at higher doses (p<0.0001). Estradiol creates weak but significant negative facilitation (p<0.05). **d**, Sensitive cells with combination treatment (n=3). Compared with single treatment with ribociclib, combination treatment with estradiol significantly reduces facilitation (p<0.0001) although completely to 0 (p=0.016). The strength of facilitation is unchanged at the lower 1nM dose of fulvestrant and increased at the higher 3nM dose (p=0.004).

1. *Facilitation will not be observed in cocultures treated with estradiol alone*. Estradiol treatment alone improves the growth of sensitive cells in monoculture, consistent with sensitive cells being more dependent on estradiol signaling. However, in coculture the observed growth of sensitive cells is, in fact, slightly less than expected if the effects of competition and estradiol were independent (**Fig. 3a, c, Supp. Fig. 8b**). This result supports the hypothesis that without selective pressure, competition from sensitive cells blocks resistant cell growth and facilitation of sensitive cells.
2. *Resistant cells will facilitate sensitive cells when treated with other agents that disrupt the ER pathway, such as the ER antagonist, fulvestrant*. In monoculture, cells resistant to ribociclib are also more resistant to fulvestrant, due to their lower dependence on estrogen signaling. In coculture treated with fulvestrant, resistant cells were found to facilitate sensitive cells, allowing them to proliferate more than expected and enough to eventually outcompete resistant cells entirely (**Fig. 3b-c, Supp. Fig. 8e**), with dose-dependent effects that parallel those with ribociclib.

Our hypothesis also makes two specific predictions about how facilitation is changed by addition of estrogen pathway modifiers. To quantify these effects, we compare observed growth with a null model that assumes that the effects of treatments are independent.

1. *Estradiol will cancel the facilitation of sensitive cells by resistant cells in the presence of ribociclib*. Addition of estradiol nearly cancels the facilitation of sensitive cells by resistant cells when treated with ribociclib (**Fig. 3a, d, Supp. Fig. 8c, f**). We see no dose response likely due to saturation of estradiol binding capacity of sensitive cells.
2. *Under ribociclib treatment, the addition of fulvestrant will strongly modify facilitation*. When ribociclib is combined with fulvestrant, facilitation of sensitive cells is unchanged at a low dose and increased at a higher dose (**Fig. 3b**), consistent with resistant cell facilitation of sensitive cell growth under treatment by each of ribociclib and fulvestrant separately (**Fig. 3d**).

Letrozole, an aromatase inhibitor (AI), only partially cancels facilitation, due to the existing resistance of CAMA-1 cells (**Supp. Fig. 9a, b**). Other endocrine inhibitors, tamoxifen (**Supp. Fig. 9c, d**) and raloxifene (**Supp. Fig. 9e, f**), both selective estrogen receptor modulators (SERM), completely cancel facilitation by disrupting the response of sensitive cells. The resulting degree of facilitation modification between the treatment with a SERM or SERD is likely due to differences in each inhibitor’s mechanism of action. While both agent classes inhibit ER signaling, they do so in different ways. SERMs bind to ER, resulting in an inactive complex, while SERDs disrupt signaling by preventing dimerization and targeting ER for degradation. In our *in vitro* system, cells resistant to ribociclib are also more resistant to the SERD, fulvestrant. However, SERMs are more damaging to resistant cell growth while also blocking facilitation in sensitive cells as they directly competing with estradiol at the receptor site, disallowing estradiol from binding to the receptor which leads to the cancellation of facilitation.

### Coculture results in a shift of sensitive cells to a more resistant cell state as revealed by scRNAseq

We hypothesized that hormone communications from resistant to sensitive cells activates proliferative pathways of sensitive cells when cocultured during ribociclib treatment resulting in a more resistant phenotype. We performed scRNAseq on sensitive and resistant CAMA-1 cells grown in 3D after 11 days of treatment, both in monoculture and coculture with initially equal proportions. Following quality control filtering and normalization of the scRNAseq profiles (see **Methods**), we performed UMAP dimension reduction to compare the phenotypic similarity of cells across the transcriptome. Monoculture sensitive and resistant lineages formed separate clusters, indicating that they are phenotypically distinct, with a shift of sensitive cells towards resistant cells seen when grown in coculture (**Fig. 4a**). We next calculated the cell cycle phase of each cell using canonical markers (**Fig. 4b**) and found a striking increase in the proportion of cycling sensitive cells in coculture compared to sensitive cells in monoculture. This analysis revealed that resistant cells facilitated the cell cycle progression of sensitive cells under drug pressure.

**Figure 4.**
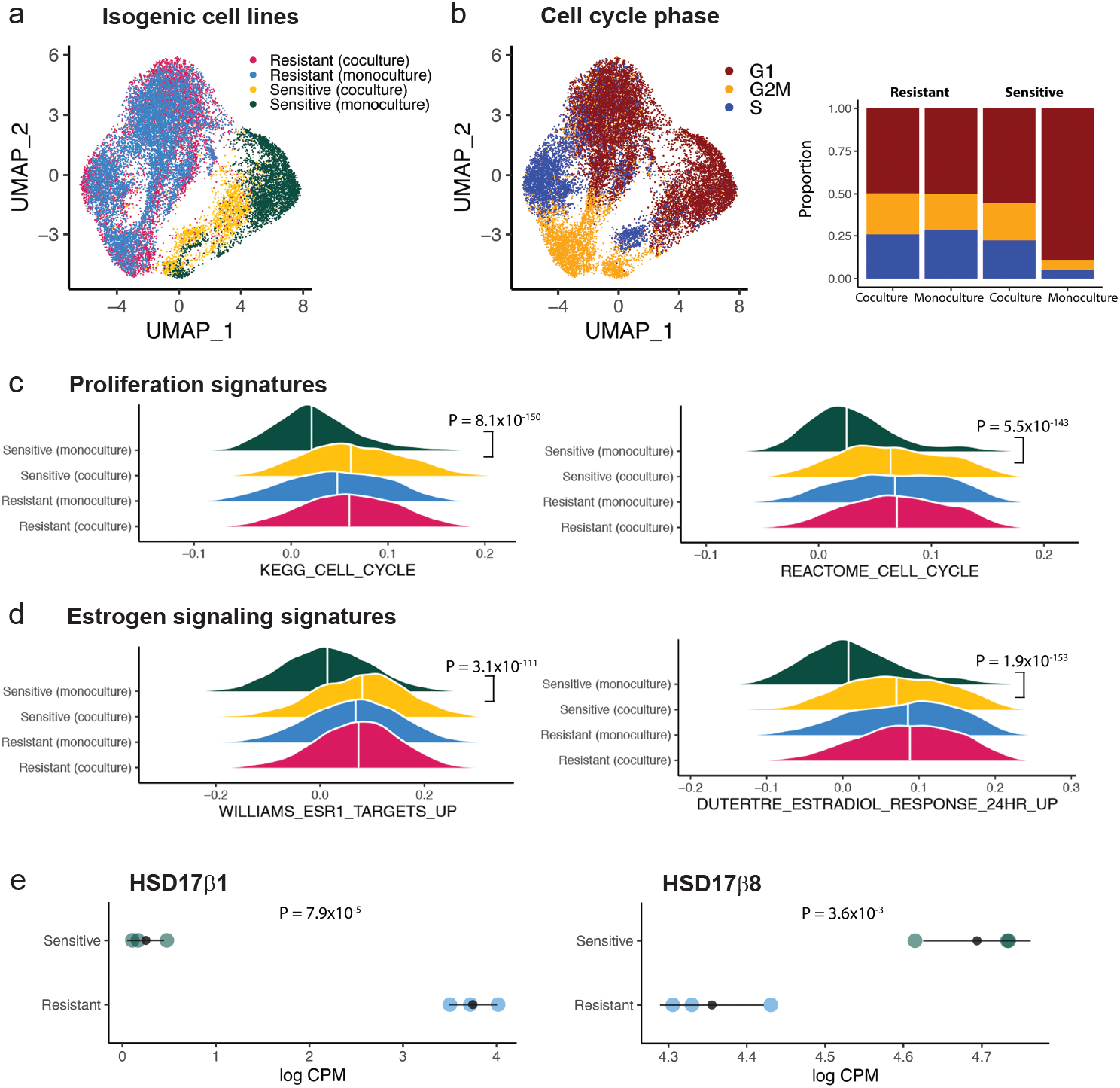
RNA sequencing reveals transcriptomic differences between mono- and cocultured sensitive and resistant cells. **a**, UMAP of scRNA-seq profiles from 6292 resistant (coculture), 7557 resistant (monoculture), 2059 sensitive (coculture) and 3329 sensitive (monoculture) CAMA-1 cells. **b**, Cell phenotypic heterogeneity visualized using UMAP and colored according to cell phase estimated using the scRNA-seq profiles of CAMA-1 cells. Stacked bar plots on the right show proportion of cells within each phase of the cell cycle. **c**, Ridge density plots showing density of ssGSEA enrichment scores for cell cycle pathways across different cell lines. White vertical line indicates median of the distribution, with FDR adjusted p-value from Wilcoxon rank-sum test between sensitive monoculture vs. sensitive coculture indicated to the right. **d**, Ridge density plots showing density of ssGSEA enrichment scores for estrogen signaling response pathways across different cell lines. White vertical line indicates median of the distribution, with FDR adjusted p-value from Wilcoxon rank-sum test between sensitive monoculture vs. sensitive coculture indicated to the right. **e**, Dot plots showing gene expression levels measured in ribociclib resistant and sensitive cells using bulk RNA-seq. Each colored dot indicates one independent biological replicate, with the grey dot and bars indicating mean and standard deviations. The p-value indicate significance of difference in means from Welch’s t-test.

We next investigated the key pathway phenotypes acquired by the sensitive cells in coculture compared to monoculture using single sample gene set enrichment analysis. We found a highly significant increase in the KEGG and REACTOME cell cycle pathway enrichment scores (both p<0.00001), supporting the cell cycle phase analysis results (**Fig. 4c**). Differential expression analysis confirmed that both proliferation and estrogen signaling gene expression was elevated in sensitive cells when cocultured compared to in monoculture (**Supp. Fig. 10**).

Gene expression signatures indicative estrogen signaling activation, including ESR1 targets (p = 3.1 × 10^−111^) and estradiol response (p=1.9 × 10^−153^), were also elevated in sensitive coculture cells (**Fig. 4d**). Finally, differential expression of estradiol production enzymes was detected between resistant and sensitive cells with increased levels of HSD17β1 in resistant cells while HSD17β8 was elevated in sensitive cells (**Fig. 4e**), with low/non-detectable aromatase expression.

### Measuring the impacts of growth factor mediated facilitation

To measure the impacts of estradiol mediated facilitation, we constructed stage-structured consumer-resource differential equation models that describe the production and uptake of estradiol, separate cell division and death, and include functional forms for the effects of ribociclib and estradiol **(Fig. 5a)** (detailed in Methods and Supp. Inf.). We used Bayesian inference to fit the model to the growth of monocultures and cocultures of sensitive and resistant cells across eight doses of ribociclib.

**Figure 5.**
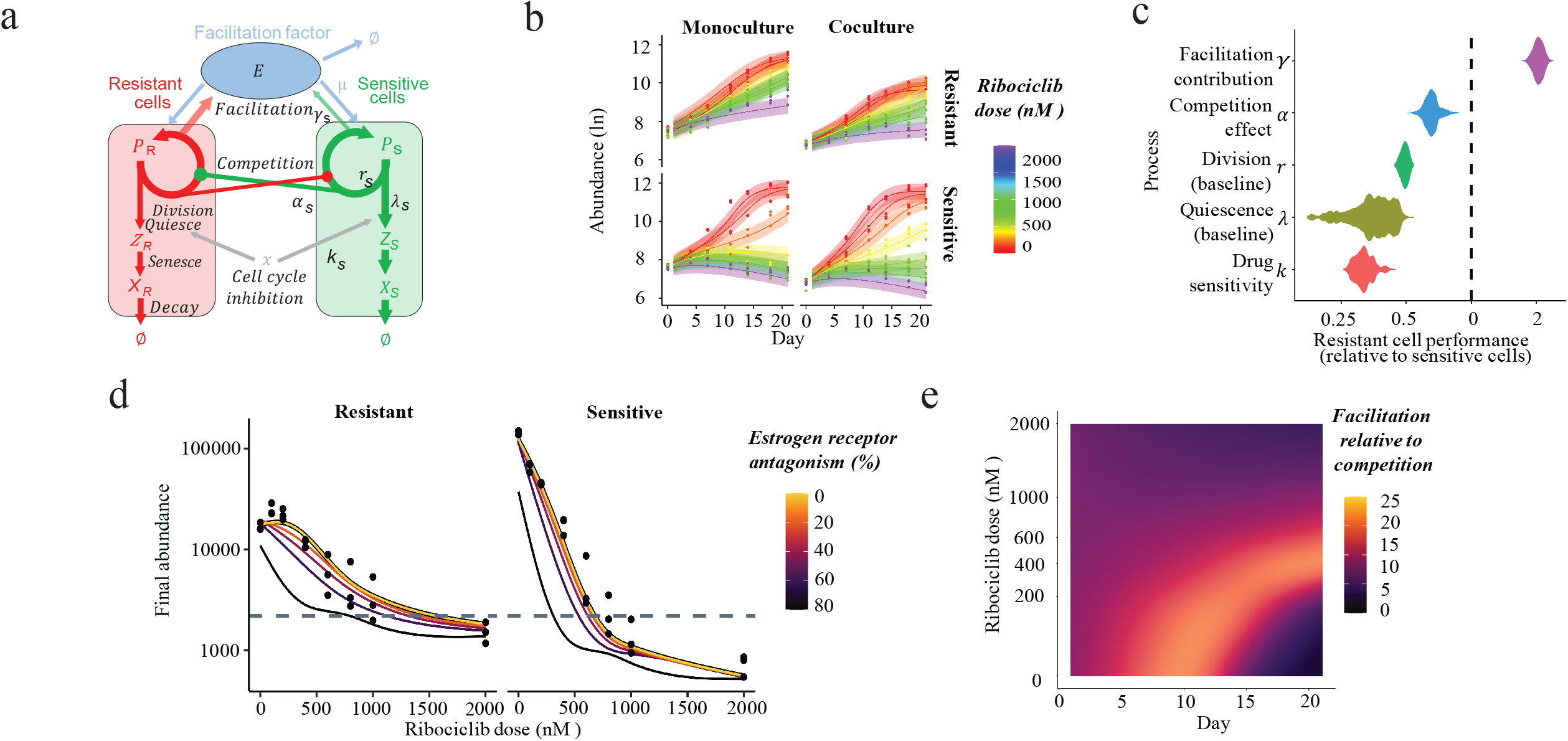
Measuring the strength and role of facilitation during treatment. **a**, Structure of mechanistic stage–structured differential equation model of facilitation fitted to data. This describes the abundance of resistant (red) and sensitive (green) cells which compete (circle headed arrows) and also facilitate one another through the production of facilitation factor (E; estradiol; blue) used by both cell types to promote cell division. Resistant and sensitive cells transition through proliferative (P), quiescent (Z) and then senescent (X) stages. Proliferative cells enter the G1/S cell cycle checkpoint at a rate that depends on competitor abundance and facilitation factor concentration. Upon entry to the G1/S cell cycle checkpoint, cells either divide or enter the quiescent state. Cell cycle inhibitor therapy prevents accumulation of cell cycle promoting proteins, increasing the fraction of cells entering the quiescent stage, rather than dividing. **b**, Predicted spheroid growth from initial abundance for sensitive and resistant cells (rows), alone or in co-culture (columns) and across a broad range of ribociclib doses (color). Observed cell counts over time (points) are accurately predicted across all doses and cell type compositions (lines) by the facilitation model and uncertainty in model fits are captured by the 95% high credibility intervals (shaded regions). **c**, Quantification of the resistant and sensitive cells’ abilities to facilitate neighboring cell growth, compete for resources, divide or quiesce in the absence of therapy and to respond to the direct impacts of treatment (drug sensitivity). Performance of resistant cells, relative to sensitive cells, is shown for each process (y axis), with the violin plots showing the range of relative performance values yielding spheroid trajectories consistent with the observed data (shaded region). Equal performance of the two cell types is signified by the vertical dashed line. **d**, Prediction of impacts of blocking facilitation on sensitive and resistant cell growth in co-culture, across drug dose. Datapoints show the observed final size of replicate spheroids across doses without facilitation blocking (100% facilitation). The thick black bordered lines show the facilitation model predictions of final size. Thinner lines show the predicted reduction in final abundance of resistant and sensitive cells achievable by reducing facilitation (brighter lines indicate greater facilitation levels). **e**, The role of facilitation on spheroid growth, relative to competition, across drug doses. Each point on the surface indicates a specific time and dose. The coloration shows how important facilitation is at that moment (brighter coloration indicates that facilitation is more important). The bright band shows the window of facilitation when abundance of both cell types is increasing but the carrying capacity (maximal density of the spheroid) is not reached.

The model of estradiol mediated facilitation accurately describes the growth of resistant, sensitive and coculture spheroids across all drug doses **(Fig. 5b)** and additionally quantifies how biological processes differed between resistant and sensitive cells **(Fig. 5c)**. Resistant cells were found to be approximately 60% as competitive (*α*), consistent with the 1.5 times higher competitive effect of sensitive cells (**Fig. 1f)**. Net production of estradiol (*γ*) by resistant cells was estimated to be approximately twice that of sensitive cells, consistent with the ratio of production/use estimated directly (8.0/2.5 = 3.2, **Fig. 2c**). In the absence of treatment, the less competitive resistant cells had a baseline division rate (*r*) 50% of sensitive cells and a lower quiescence rate (*λ*). However, they were far less sensitive to the cell cycle inhibitory effects of ribociclib (*k*), allowing their spheroids to continue to grow at high doses.

We verified that models of alternative mechanisms of cell interaction could not produce the diversity of spheroid growth trajectories observed across drug doses and compositions. Alternative models included direct competition for resources and phenotypic plasticity in which cells transition from a naive to a resistant state either in response to drug induction or via random switching (detailed model comparisons provided in Supp. Inf.). We performed formal probabilistic model comparison of the non-nested set of model hypotheses using Watanabe–Akaike information criterion (WAIC). The facilitation model greatly outperformed models of competition or phenotypic plasticity (**Supp. Fig. 11)**. This analysis guarded against model overfitting and supports the estradiol mediated facilitation hypothesis.

We used the estradiol mediated facilitation model to explore the impact of therapeutically blocking facilitation on the growth of sensitive and resistant co-cultured populations. In the model, we reduced the binding of estradiol to its cellular receptor (*μ*) by differing degrees to reflect exposure to different doses of a receptor blocking drug such as fulvestrant. In the absence of facilitation blocking, the facilitation model accurately predicts the final resistant and sensitive cell population size (**Fig. 5d**; points show observed final cell counts at day 21, black line + yellow overlay shows model predictions under the experimental conditions). However, facilitation needed to be reduced by more than 50% before substantial reductions in spheroid growth were predicted, suggesting that resistant cells produce sufficiently large amounts of estradiol to saturate the growth benefits of facilitation. At lower ribociclib doses (<500nM), facilitation blocking caused a more severe reduction in the abundance of sensitive than resistant cells, suggesting that resistance could be promoted under these circumstances.

Finally, we assessed the importance of facilitation relative to resource competition in determining the rate of spheroid growth throughout treatment **(Fig. 5e)**. Across ribociclib doses (<600nM) we showed that facilitation was many times more important than competition during the period after the accumulation of a sufficient population size to generate high levels of estradiol but prior to the spheroid reaching maximal density and becoming regulated by resource limitation. At moderate ribociclib doses (200-600nM), the facilitation window shifts later into the treatment, as resistant cell abundances increase more slowly, and the window also extends for a longer duration. At the highest ribociclib doses (>600nM), the window of facilitation closes completely because proliferation of resistant facilitating cells is controlled (shown in **Fig. 5e**). Coculture spheroid size then shrinks during treatment as the small resistant population alone cannot maintain estradiol levels to overcome treatment.

## Discussion

We developed *in vitro* model systems to investigate how interactions between cancer lineages impact the growth of heterogeneous ER+ breast cancer populations. We found that cells sensitive to ribociclib (CDK 4/6 cell cycle inhibitor) grow faster in untreated monoculture and outcompete resistant cells in coculture. However, in the presence of ribociclib, resistant cells facilitate the growth of sensitive cells by producing estradiol, a potent estrogen metabolite, and upregulation of estrogen signaling and proliferation. Although resistant cells were developed by long-term growth with ribociclib, they also acquired cross-resistance to endocrine therapy due to a common signaling pathway and thus also facilitate growth of sensitive cell populations in the presence of ER antagonists. These patterns of cross-resistance and cell-cell interactions complicate patient treatment even beyond the challenge of evolution of resistance.

We use mathematical models to quantify competitive and facilitative interactions, and to predict how different conditions will alter facilitation. Mathematical models that include facilitation improve our ability to describe the growth of heterogeneous cancer populations. These models predict that blocking facilitation has the potential to control both resistant and sensitive cell populations and inhibit the emergence of a refractory population, prolonging the benefits of ribociclib. This may explain why ribociclib is more effective when combined with antiestrogen therapy than as monotherapy for some patients (38-40).

Our dynamical models highlight that the contribution of facilitation to proliferation and cancer population growth shifts during the growth of the cancer. As cell abundance increases, facilitation-driven growth diminishes as carrying capacity is reached, due to an increase in resource competition. This shift suggests that competitive ability may be more essential at later stages in disease progression, while facilitation may be more important in earlier stages. Future studies may also integrate local spatial effects in our measurements and modeling to account for any impact in cell-cell interactions.

Mathematical modeling (41-43) and experimental (44-46) research proposes that tumor growth and progression may be promoted by cooperation among diverse cell populations by sharing of resources or products through mechanisms including neoangiogenesis (41) and growth factor production (46). Our results support the concept that cancer cell facilitation may impact heterogeneous tumor growth during treatment in ER-positive breast cancer. No current therapy is directed specifically at cooperative cancer cell interactions. However, as tumors are often comprised of multiple subclonal populations, interactions between cells may provide targets for long-term treatment strategies. Given that facilitation can increase both persistence and biomass relative to simpler systems with resource competition only (47, 48), it is likely to be frequent in heterogeneous cancer populations. As we gain in capabilities to measure the growth of heterogeneous subclone populations during treatment, we may tailor strategies to control faster growing yet drug sensitive cells while preventing dominance of refractory cells through competition with more fit subclones.

This study describes facilitation through an increase of local estradiol concentrations by resistant cells that stimulates proliferation of sensitive cells during selective pressure. It is known that estrogen biosynthesis and metabolism can be upregulated in cancer cells (49-51). It remains unknown whether resistant cells in our system acquired mutations to increase estradiol levels or if changes are epigenetic. The increase of active estrogen is important given that the majority of breast cancers are hormone receptor positive, and resistance to first line standard of care hormone therapy is common (52-54). In patients with increased levels of estradiol, potential treatment strategies include blocking the enzymes that synthesize estradiol in resistant cells or modulating the level or type of endocrine therapy given to a patient. Targeting the diffusible facilitation factor has been proposed to be a more evolutionarily stable treatment strategy than targeting the cancer cell receptors or signaling pathways (55) because the production provides little benefit to the producing cell. An important next step is to study these mechanisms in patient tumors, and to better understand the presence and impact of cell cooperation in response to therapy and outcomes.

In conclusion, our data indicate that the facilitation within heterogeneous cancer cell populations shapes the dynamics of resistance and cancer growth of the population as a whole. We show that local production of a highly active estrogen, estradiol, facilitates the growth of sensitive cells in the presence of drug treatment, is produced by resistant cells, and conveys resistance to anti-proliferative effects of cell cycle inhibition in sensitive cells. We use mathematical models to measure these processes and show that blocking this facilitation could promote response of the entire cancer population, reducing selection for refractory resistant states. This work provides support for the development of strategies to modulate facilitation to create robust and durable cancer control.

## Methods

### Cell lines and reagents

The previously authenticated estrogen-receptor-positive (ER+) CAMA-1 breast cancer cell lines were maintained in DMEM+ 10% FBS+ 1% antibiotic–antimycotic solution. The ribociclib-resistant CAMA-1 cell line creation (ribociclib-resistant CAMA-1) was previously reported (32). Briefly, cells were cultured and continuously treated with ribociclib (Selleck Chemicals, Cat. No: S7440) at 1µM for 1 month. Following the initial 1µM ribociclib treatment, cells were treated with 250nM for 4 months to develop resistance. Maintenance of ribociclib-resistant CAMA-1 cells continued in complete culture medium + 250 nM ribociclib. Resistance against ribociclib was detected by the alteration of the dose–response curve measured using CellTiterGlo Chemiluminescent Kit (Promega Corporation, Cat. No.: G7573).

### Lentiviral labeling of sensitive and resistant cells

Using lentiviruses incorporating distinct fluorescent proteins, we labeled CAMA-1 parental sensitive cells (venus; LeGO-V2) and ribociclib resistant cells (cerulean; LeGO-Cer2). LeGO-V2 and LeGO-Cer2 vectors were provided by Boris Fehse (Addgene plasmids #27340 and #27338). Lentiviruses with fluorescent proteins were created using Lipofectamine 3000 reagent (Thermo Fisher Scientific) following manufacturer’s protocol. CAMA-1 sensitive and resistant cell lines were transduced with lentivirus using reverse transduction. Briefly, 1mL of polybrene-containing cell suspension of 200,000 cells were plated in a well of a 6-well plate. Previously, 0.5 mL of viral aliquot had been dispensed in plate. Following 48 hours of incubation at 37 °C with 5% CO_2_, cells were washed and given fresh regular culture medium. To select for fluorescence-activated cells, fluorescently labeled cells were flow-sorted after further subculture of transduced cells attain homogenously labeled cell populations.

### Mono- and coculture 3D spheroid experiments

18-21 day-long experiments were initiated with fluorescently labeled sensitive and resistant cell lines in different compositions. For CAMA-1 spheroid experiments, as earlier reported (32), 2000 cells were plated in different proportions (100% CAMA-1 sensitive, 50% CAMA-1 sensitive— 50% CAMA-1 resistant, 100% CAMA-1 resistant) in 96-well round-bottom ultra-low attachment spheroid microplate (Corning, Cat. No.: 4520). 24 h later, spheroids were washed and fresh medium including treatment drugs was applied. Spheroids were treated for a total of 18-21 days with imaging and media change performed at every 4th and 7th day of the week. Spheroids were treated with the following drug therapies at specified doses described in results and figures 1 and 4: ribociclib (Selleck Chemicals, Cat. No: S7440), estradiol (Peprotech, Cat. No: 5022822), fulvestrant (Selleck Chemicals, Cat. No: S1191). Imaging was performed using Cytation 5 imager (Biotek Instruments) recording signal intensity from brightfield, YFP (for Venus fluorescence) and CFP 450/440 (for Cerulean fluorescence) channels. Raw data processing and image analysis were performed using Gen5 3.05 and 3.10 software (Biotek Instruments). Briefly, the stitching of 2 × 2 montage images and Z-projection of 6 layers using focus stacking was performed on raw images followed by spheroid area analysis. To quantify growth under these conditions, we measured fluorescence intensity and growth of spheroid area over the total time of the experiment. For cell count calculations, a standard curve was created by measuring the area of spheroids after plating at different cell numbers 24 hours after plating. A resulting equation by fitting a curve to the data was performed by GraphPad Prism 7.02 software (second order polynomial – quadratic – curve fit used). Whole spheroid area and fluorescence intensity measurements of each population were integrated into the fitted equation, and cell counts for each population were produced from fluorescence intensities relative to spheroid size. All coculture experiments were performed in triplicates.

### Cell number quantification

For CAMA-1 cells, cell numbers were quantified as described previously (Grolmusz et al, 2020). In brief, the relationship of area to cell counts follows a nonlinear curve (**Supp. Fig. 1a**), and sensitive and resistant CAMA-1 cells have similar relationships of area to fluorescence (**Supp. Fig. 1b**). Cell numbers were estimated by inverting the non-linear function, with proportions of the two cell types estimated by the normalized relative fluorescence of each wavelength.

To estimate cell numbers for MCF7/LY2 cells, we measured cell numbers and spheroid area for a range of initial fractions of sensitive (S) and resistant (R) cells and fit to a Michaelis-Menten function maximum value A and half-saturation constant K (**Supp. Fig. 1c**). The relationship differs depending on the proportion of S cells. Parameters are K=2.753×10^5^, A=1.094×10^7^ with pure R cells and K=2.119×10^5^ and A=5.459×10^6^ with pure S cells, and K=2.330×10^5^ and A=7.546e6 ×10^5^ with mixed cultures (the dashed red curve).

To correct for possible differences in per cell fluorescence, we regressed the fluorescence of pure cultures against the known numbers of S and R cells (**Supp. Fig. 1d**). We find a slope of 8.04 ×10^4^ for S cells and 1.349×10^5^ for R cells, and thus estimate that each R cell produces 1.677 times as much fluorescence. To find the numbers of S and R cells in coculture from the area and the fluorescence, we find the total cells by inverting the relationship between cells and area, and the proportion of each cell type from the fraction of fluorescence, with R cells reduced by the weighting factor of 1.677.

To estimate cell numbers for MCF7 cells, we followed the same procedure, finding Michaelis-Menten fits (**Supp. Fig. 1e**) with parameters K=1.772×10^5^, A=9.061×10^6^ with pure R cells and K=5.315×10^5^ and A=1.138×10^7^ when there are any S cells in the culture. We correct for fluorescence in the same way (**Supp. Fig. 1f**), finding a slope of 8.36×10^4^ for S cells and 2.029×10^5^ for R cells, and thus estimate that each R cell produces 2.426 times as much fluorescence.

### Liquid chromatography-tandem mass spectrometry (LC-MS/MS)

Media samples taken at day 21 from 3D spheroid experiments, treated with or without ribociclib, (experimental setup previously described in results and methods - ***Mono- and coculture 3D spheroid experiments***) and plated in different compositions (100% sensitive, 50% sensitive – 50% resistant, and 100% resistant) were spun down at 300g and frozen at -80 °C. Samples were then prepared by the Analytical Pharmacology Core of City of Hope National Medical Center for LC-MS/MS for estrone and estradiol detection. LC-MS/MS system consisted of a Shimadzu Prominence HPLC system interfaced to an AB SCIEX QTRAP® 5500 system (Foster City, CA, USA). HPLC separation was achieved using a XSELECT CSH Phenyl-Hexyl 3.5um, 2.1×150mm analytical column (Waters). The column temperature was maintained at 50°C, and the flow rate was 0.38ml/min. The mobile phase consisted of A (Water: 1000ml + 60ul 30%NH4OH) and B (Methanol : 1000ml+ 60ul 30% NH4OH). The following gradient program was used: 55% B (0.01 min), 70% B (0.01-4.0 min), 100% B (5.5min), 30%B (8.5min). The total run time was 8.5min. The auto-injector temperature was maintained at 15°C. The atmospheric pressure chemical ionization (APCI) source of the mass spectrometer was operated in negative ion mode with ion source gas (55), curtain gas (20), collision gas (High), nebulizer current -4.0, The entrance potential was set to -10V. Declustering potential (DP) was -110, collision energy (CE), and collision cell exit potential (CXP) was optimized to -50V, -21V for Estrone, -160V, -50V,-17V for Estrone internal standard), -210V,-58V,-19V for Estradiol and -205V,-52V, -13V for Estradiol internal standard respectively. The source temperature was 400 °C. A solvent delay program was used from 0 to 3.5 min and from 6.5 to 8.5min to minimize the mobile phase flow to the source. Analyst software version 1.5.1 was used for data acquisition and processing. Atmospheric – pressure chemical ionization of Estrone, Estrone D4, Estradiol, Estradiol D5 produced abundant protonated molecular ions (MH-) at m/z 268.980, 272.983, 270.969, and 275.981 respectively. Fragmentation of these compounds was induced under collision induced dissociation conditions. The precursor→product ion combinations at m/z 268.980→145.200 for Estrone, and 272.983→147.100 for Estrone-I.S. 270.969→182.800 and 275.981→147.000 for Estradiol and Estradiol-IS were used in multiple-reaction monitoring mode for quantitation. Under optimized assay conditions, the retention times for Estrone, Estrone IS and Estradiol, Estradiol -IS. were 4.89, 4.60 min, respectively.

### Single cell RNA Sequencing

Spheroids of different compositions (100% sensitive, 50% sensitive – 50% resistant, 100% resistant) were initiated from Venus-labeled and mCherry-labeled CAMA-1_ribociclib_resistant cells and were subjected to 1 uM ribociclib treatment. After 11 days, spheroids were harvested, washed and cell suspensions were viably frozen for further processing. Once thawed, cells were centrifuged at 300 x g and washed twice with 37°C pre-warmed 1x PBS, pH 7.4 (Gibco, Cat #10010) + 0.04% Nuclease-Free Bovine Serum Albumin (BSA, EMD Millipore, Cat # 12661525mL). Cells were resuspended to a target concentration of 1,000 cell/µL, concentrations were confirmed using trypan blue staining and counted on a hemocytometer. scRNA-Seq was performed on resuspended cells using the 10X Genomics Chromium Single Cell 3’ GEM, Library, & Gel Bead Kit v3 (10X Genomics, Cat # 1000075) according to manufacturer instructions at a target of 10,000 cells per sample. Each sample was barcoded with a unique i7 Index during library preparation using the Chromium i7 Multiplex Kit (10X Genomics, Cat# 120262) to allow sample multiplexing during sequencing. Libraries were sequenced at Fulgent Genetics, on an Illumina HiSeq X Ten instrument with 2×150 paired end reads, to a read depth of 10,000 reads per cell; additional sequencing was performed on an Illumina NovaSeq 6000 with 2×150 paired end reads increase read depth to a total depth of 50,000 reads per cell.

### Single Cell Analysis

scRNA-Seq was performed at the Integrative Genomics Core at City of Hope, Fulgent Genetics, and the High Throughput Genomics Core at Huntsman Cancer Institute (HCI) of University of Utah. Sequence reads were processed with CellRanger v3.0.2 using reference genome (GRChg37). A gene-barcode matrix was generated for each sample containing counts of unique molecular identifiers (UMIs) for each gene in each barcode (cell). The matrix was processed with Seurat v3.1.1.9023 (56) to identify cell populations. A series of filters were applied to the data before performing clustering. First, high-quality cells were retained by using the following filters in Seurat: subset = nFeature_RNA < 7000 & nFeature_RNA > 3000 & nCount_RNA > 2000 & nCount_RNA < 60000 & percent.mt < 30. Second, doublet cells were predicted with scrublet (threshold=0.25) (57) and the predicted doublets were removed from further analysis. Third, cells expressing mCherry or mVenus were retained by using cells having 2 or more UMI counts of either mCherry or mVenus. The filtered UMI count matrix was normalized with method “LogNormalize” and “scale.factor=10000”. The top variable 2000 genes were identified and were used to perform Principal component analysis (PCA) and clustering in Seurat. Cell clusters were visualized using Uniform Manifold Approximation and Projection (UMAP). Differential expressed genes between cell populations were identified using MAST (58) as implemented in Seurat using FindMarkers function (abs(foldchange) >= 0.2 & Adjust P value <= 0.05). Genes were ranked based on fold change. Pathway analyses were performed on 50 hallmark signatures (MSigDB, hallmark) (59) and 4725 curated pathway signatures (MSigDB, c2) using single samples gene set enrichment analysis implemented in the R package GSVA (60). Significant differential pathway activity was identified using Wilcox rank sum test.

### Western blot analysis of CAMA-1 cells

Lysates of CAMA-1 cells were separated by SDS-polyacrylamide gel electrophoresis and proteins were transferred electrophoretically to a polyvinylidene difluoride membrane using Invitrogen iBlot 2 device and Invitrogen iBlot Transfer Stacks. Membranes were blocked with Tris-buffered saline with 0.05% tween 20 (TTBS) and 5% BSA for 1 hour at room temperature. After washing with TTBS, membranes were then probed with anti-HSD17β1 polyclonal antibody (Abnova, H00003292-M03A, 1:1000 dilution, overnight 4°C; R&D systems, MAB7178, 1:2000 dilution, overnight 4°C), anti-HSD17β8 polyclonal antibody (Proteintech, 16752-1-AP, 1:1000 dilution, overnight 4°C), and anti-β-actin monoclonal antibody (Santa Cruz Biotechnology, sc-47778, 1:500 dilution, 1 hour room temperature) and detected using SuperSignal West Pico PLUS Chemiluminescent Substrate (Thermo Scientific) with anti-rabbit (GE Healthcare NA9341ML) or anti-mouse (GE Healthcare NXA9311ML) peroxidase-linked secondary antibody (1:6000 dilution). The molecular weight was determined using a pre-stained protein marker (BioRad).

### Estradiol production and uptake analysis

To estimate the production and use of estradiol by sensitive and resistant cells, we compare the fits of three functions, each based on Michaelis-Menten kinetics by minimizing the least squares difference from the observed estradiol concentration. The full model includes separate production and uptake rates for each cell type. We estimate the production parameters *ρ*_*R*_ and *ρ*_*S*_ and the uptake parameters *α*_*R*_ and *α*_*S*_, where *R* and *S* represent the measured cell numbers, with the function:

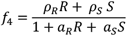

We examined simpler models that restrict the values. To test a model without production by sensitive cells, we set *ρ*_*S*_ = 0, giving:

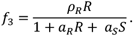

Because uptake by cells is large compared with background usage (scaled to 1 in each of these models), we tested a simplified model with background usage set to zero. Without that term, the model is overparameterized, and we scale *α*_*R*_ = 1. In this case, *αα*_*SS*_ represents per cell uptake by sensitive cells relative to resistant cells, and the function simplifies to:

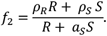

During model fitting, we eliminated one outlier (Sample F8). We used the reported lower limit of detection of LLOD = 62.5 and set all measured values of 0 to LLOD/2. Neglecting this adjustment leads to significantly worse fits.

Model *f*_3_ provided a poor fit. The best fit with model *f*_4_ estimates *ρ*_*R*_ = 83.58, *ρ*_*S*_ = 10.11, *α*_*R*_ = 0.864, *α*_*S*_ = 0.345 and a residual sum of squares of 7321.1. The best fit parameters with model *ff*_2_ have the same residual sum of squares, with estimates *ρ*_*R*_ = 96.79, *ρ*_*S*_ = 11.71, *α*_*S*_ = 0.400. Due to the equally good fit with one fewer parameter, we choose this as our final model.

To find confidence limits, we use the residual sum of squares to estimate the variance and convert the least squares into a likelihood. We find values of each parameter with log likelihood within 2 of the maximum. With model *f*_2_, we find limits *ρ*_*R*_ ∈ [91.49,102.08], *ρ*_*S*_ ∈ [9.88,13.54], *α*_*S*_ ∈ [0.355,0.460].

### Mathematical modeling – FACT analysis

#### Facilitation Analysis through Combination Therapy

The FACT algorithm breaks into six steps, which we outline here before describing in detail below. Modifiers refer to estrogen pathway modifying treatments, including estradiol itself and three ER antagonists: fulvestrant, tamoxifen, and raloxifene.

1. **Fit growth and carrying capacity** for each cell type in monoculture with no treatment.
2. **Treatment effects:** Using the carrying capacity in the absence of treatment, find the effects of ribociclib or modifier treatment on cell growth and death rates.
3. **Synergy:** From growth with combined ribociclib and modifier treatment, compare observed growth with that expected under a null model without interaction.
4. **Competitive effect:** Using the carrying capacities and growth in sensitive and resistant cells in monoculture from Step 1, estimate competition coefficients between the two cell types in coculture.
5. **Facilitation:** Using the carrying capacities, growth and death rates of single cell types with or without ribociclib from Steps 1 and 2, and the competition coefficients in the absence of treatment from Step 3, quantify facilitation as deviations of growth from predicted under a null model. To quantify facilitation in the presence of modifier we follow the same steps but with modifier treatment.
6. **Facilitation modification:** Using the growth of cells in competition with ribociclib (Step 5) and the direct effects of modifier (Step 2), quantify whether the modifier enhances or reduces facilitation.

### Mathematical methods

We fit each model with least squares, using all three replicates and the observed mean on day 4 for the initial condition. We exclude the measurements before day 4 because the cells often show a lag before beginning their growth.

#### 1. Fit growth and carrying capacity to single cell data in the absence of treatment

We use the logistic model, as defined by the differential equations for sensitive and resistant cells

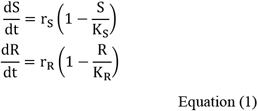

to estimate *r*_*S*_, *r*_*R*_, *K*_*S*_ and *K*_*R*_ as the growth rates and carrying capacities of sensitive (S) cells and resistant (R) cells.

#### 2. Treatment and modifier effects

Using the carrying capacity in the absence of treatment we estimate growth with treatment (*r*_*ST*_, *r*_*RT*_) as shown, and with modifier (*r*_*SM*_, *r*_*RM*_) with the model

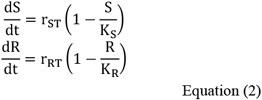

#### 3. Synergy

To quantify how ribociclib interacts with modifiers in monoculture, we fit the logistic model using carrying capacities from Step 1, and compare the estimated growth *r*_*C*_ with the null model:

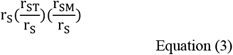

#### 4. Estimate competition coefficients

Using the estimates of the growth rates and carrying capacities from Step 1, we fit untreated coculture data to a Lotka-Volterra competition model with competition coefficients *α*_*SR*_ and *α*_*RS*_

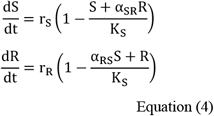

#### 5. Estimate facilitation

We estimate facilitation as the deviation between observed and expected growth. To find expected growth, we use the estimates of the growth rates in monoculture with treatment, carrying capacities from the untreated monoculture, and the competition coefficients from the untreated cocultures.

#### 6. Facilitation modification

If the presence of the other cell type enhances growth in the presence of ribociclib, a modifier could cancel or enhance this effect. To quantify, we use the growth parameter from step 4 in the presence of ribociclib, and account for the direct effects of the modifier as in step 2.

### Facilitation Analysis by Combination Therapy – Mechanistic model

#### Facilitation factor production, flux and utilization

To describe the mechanisms of facilitation between resistant (R) and sensitive (S) cells, we model the intracellular concentration of the facilitation factor (E) in each cell type (E_S_, E_R_) and the flux of this factor between cell types via the shared extra-cellular environment (E_E_). We describe the intracellular production of this factor by sensitive and resistant cells at rate ρ_S_ and ρ_R_ and the receptor binding for utilization as a growth promoting signal at rate μ_S_ and μ_R_. Facilitation factors diffuse between cells types and the extra-cellular environment at rate η. Finally, we describe the influx (σ_E_) and decay (δ_E_) of facilitation factors from the external environment. This leads to the following set of differential equations describing cellular and environmental concentrations of facilitation factors:

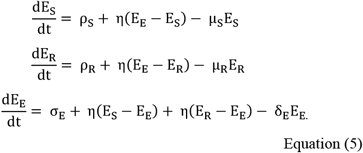

The equilibrium of this system is:

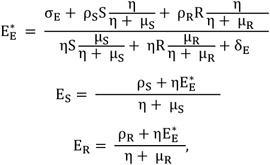

showing that the intra- and extracellular concentrations depend on the net balance between sources and sinks of facilitation factors. Inhibitors of the production of facilitation factors will reduce the intra and extracellular inputs to E (σ and ρ). In contrast drugs targeting facilitation factor receptors will reduce the binding of intracellular factors, reducing μ by some factor.

### Coupling facilitation factor dynamics with cancer coculture spheroid growth

The cellular level description of facilitation was next scaled to describe the populations of resistant and sensitive cells competing and facilitating in coculture spheroids over time. The resistant and sensitive cells are described as transitioning between proliferative (P), quiescent (Z) and senescent (X) states.

Resource competition between all cells is described by 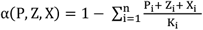, where competitive ability of each cell type (i) is determined by the carrying capacity parameter K_i_.

Proliferative cells enter the G1/S phase cell cycle checkpoint at a baseline rate (r), which is reduced by resource competition and increased by facilitation factor availability (E). This beneficial facilitation factor effect saturates at high concentrations when uptake and binding becomes rate limited (c). The proliferation rate of resistant and sensitive cells will be related to the binding of intracellularly available facilitation factors (μ_i_E_i_). The G1/S phase entry of cells of type i, based on competition and the internal concentration of facilitation factors therefore follows:

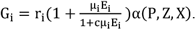

After entering the cell cycle checkpoint, cells undertake the decision to divide or enter a quiescent state, based on the balance of key regulatory cell cycle promoters and inhibitors. In addition to a baseline quiescence rate (λ_i_), cells are promoted to enter the quiescent state by the cell cycle inhibitor ribociclib (x), which inactivates the key regulators of the G1/S phase cell cycle checkpoint (CDK4/6), blocking cell cycle progression. We describe the effect of ribociclib in inhibiting cell cycle progression and driving cells into a quiescent state as increasing with drug concentration following:

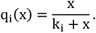

No additional quiescence is induced in the absence of treatment, complete cell cycle arrest is achieved at very high levels of ribociclib and the half maximal cell cycle arrest is achieved at a dose k_i_for cell type i. Differences in this parameter between resistant and sensitive cells describes innate resistance to the drug that emerged through selection. Following quiescence, we describe the transition of cells into a senescent state at rate φ_i_ before cell death occurs at rate δ_i_.

Combining these models of competition, facilitation, cell cycle progression, arrest and cell death yields the following spheroid population model, describing the abundance of proliferative quiescent and senescent resistant and sensitive cells in coculture spheroids over time. The dynamics of the extracellular facilitation factor concentration in the spheroid population model is governed by the balance between net secretion by proliferative and quiescent cells (resistant= γ_R_ and sensitive= γ_S_) and its decay (δ_E_), giving:

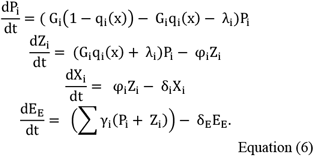

### Resistant and sensitive cell coculture spheroid growth and facilitation across drug doses

Resistant and sensitive cells were fluorescently labeled as previously described in methods (***Lentiviral labeling of sensitive and resistant cells***) and replicate populations (n=3) were plated either in monoculture or in 50:50 initial coculture. Cancer spheroids were grown in 3D culture for 21 days with imaging performed at 3-4 day intervals and the abundance of each cell type enumerated as described previously in methods (***Mono- and coculture spheroid experiments***). To explore the consequence of cell-cell interactions of drug response, the replicated mono- and coculture time course experiments were conducted under 8 ribociclib concentrations within the EC20-50 range (0,100,200,400,600,800,1000,2000nM). Spheroid analysis and cell counts performed as described previously in methods (***Mono- and coculture spheroid experiments***).

### Measuring the strength of competitive and facilitative interactions between subclones and vital cellular rates

We measured the strength of competitive and facilitative interactions between subclones, the drug impacts on cell cycle arrest and the vital cellular rates of proliferation, quiescence and senescence, by fitting the spheroid population model to the experimental data described above. Given the initial spheroid size and composition and the ribociclib concentration, the spheroid population model projects the abundance of sensitive and resistant cells throughout the experiment based on a given combination of biological rates. A log-normal distribution measurement model was used to evaluate the likelihood of each spheroid abundance observation and weakly informative Bayesian priors were used in all cases (Supp. methods). The biological rates of each process in the spheroid population model were inferred with uncertainty, using Bayesian Hamiltonian Monte Carlo (HMC) (61). Biological rates were identified that yield the most accurate model predictions of the observed spheroid growth trajectories and composition over time and across all drug doses. The HMC algorithm efficiently samples the posterior distribution, describing the likely biological rates given the model, using derivatives of the probability density function. As a result, the inference approach jointly and efficiently: i) identifies the most probable range of subclone interaction strengths (competition and facilitation), ii) predicts the cellular composition and estradiol concentration throughout the experiments and iii) provides a probabilistic measurement of the likelihood of the hypothesis encoded by the model given the available data.

### Model predictions of final spheroid size when modulating the strength of facilitation

To explore the consequences of blocking facilitation, we analyzed the impact on resistant and sensitive cell abundances of reducing the rate of receptor binding with facilitation factors (e.g. estradiol). The most likely values of the receptor binding parameters of each cell type (μ_S_ and μ_R_) were reduced by a factor ζ (between 0 and 80% reduction). Using the model and holding all other parameters constant, the predicted abundance of sensitive and resistant cells after 21 days of coculture was forecasted, assuming continuous facilitation blocking throughout this period.

## Acknowledgements

The authors would like to acknowledge the professional support of the Analytical Pharmacology Core of City of Hope National Medical Center, led by Dr. Tim Synold. The authors also thank Dr. Adam Cohen, Dr. Benjamin Decato, Dr. Jeffrey Chang, Dr. Phillip Moos, and Patrick Cosgrove for their comments on the manuscript. RSS was supported by the CSBC/PS-ON Summer Undergraduate Research Program. This research and JIG, FRA, and AHB have been supported by an NIH NCI U54CA209978 and 1U01CA264620-01 grants from the National Cancer Institute.

## Author contributions

RE carried out experiments, was involved in their analysis and wrote the manuscript. JIG and FRA carried out the mathematical modeling and facilitation analysis and wrote the manuscript. VKG generated cell lines, performed experiments, and developed system for growth measurements. AHB conceived the study, was involved in data analysis, and wrote the manuscript. RSS developed the preliminary mathematical models. AN, JC, and EFM carried out RNA sequencing analysis. All authors approved the final version of the manuscript.

## Competing interests

The authors declare no competing interests.

## Data availability

All data involving results and conclusions are provided as supplemental data with this manuscript.

## Notes

### Competing Interest Statement

The authors have declared no competing interest.

### Summary of Updates

Main figures and results revised; supplemental files updated.

