## Supplemental Information for "Ecological interactions in breast cancer: Cell facilitation promotes growth and survival under drug pressure"

### Methods

#### ***Cell lines and reagents***

The previously authenticated estrogen-receptor-positive (ER+) LY2 and MCF7 breast cancer cell lines were maintained in DMEM+ 10% FBS+ 1% antibiotic–antimycotic solution. Ribociclib-resistant LY2 and MCF7 cell lines were created. Briefly, cells were cultured and continuously treated with ribociclib (Selleck Chemicals, Cat. No: S7440) at 1 µM for 1 month for LY2 cells and 7 weeks for MCF7 cells. Following the initial 1 µM ribociclib treatment, LY2 cells were then treated with 2 uM for 4 months while MCF7 cells were treated with 1.5 µM ribociclib for 3 months to develop resistance. Maintenance of ribociclib-resistant LY-2 cells continued in complete culture medium + 2 uM ribociclib. Maintenance of ribociclib-resistant MCF7 cells continued in complete culture medium + 1.5 µM ribociclib. Resistance against ribociclib was detected by the alteration of the dose–response curve measured using CellTiterGlo Chemoluminescent Kit (Promega Corporation, Cat. No.: G7573).

#### ***Lentiviral labeling of sensitive and resistant cells***

Ribociclib resistant lines were evolved under continuous exposure to treatment and with acquired resistance confirmed by dose-response experiments (as described in methods – ***Cell lines and reagents***). Using lentiviruses incorporating distinct fluorescent proteins, we labeled parental sensitive cells (venus; LeGO-V2), ribociclib resistant LY2 cells (cerulean; LeGO-Cer2), and ribociclib resistant MCF7 cells (mCherry; LeGO-C2). LeGO-V2, LeGO-Cer2, and LeGO-C2 vectors were provided by Boris Fehse (Addgene plasmids #27340, #27338, and #27339). Lentiviruses with fluorescent proteins were created using Lipofectamine 3000 reagent (Thermo Fisher Scientific) following manufacturer's protocol. LY2 and MCF7 sensitive and resistant cell lines were transduced with lentivirus using reverse transduction. Briefly, 1mL of polybrene-containing cell suspension of 200,000 cells were plated in a well of a 6-well plate. Previously, 0.5 mL of viral aliquot had been dispensed in plate. Following 48 hours of incubation at 37 $^{o}$C with 5% $CO_2$, cells were washed and given fresh regular culture medium. To select for fluorescence-activated cells, fluorescently labeled cells were sorted after further subculture of transduced cells attain homogenously labeled cell populations.

#### ***Mono- and coculture 3D spheroid experiments***

18-21 day-long experiments were initiated with fluorescently labeled sensitive and resistant cell lines in different compositions. For LY2 spheroid experiments, 5000 cells were plated in different proportions (100% LY2 sensitive, 80% LY2 sensitive —20% LY2 resistant, 100% LY2 resistant) in 96-well round-bottom ultra-low attachment spheroid microplate (Corning, Cat. No.: 4520) while 5000 cells were plated in different proportions (100% MCF7 sensitive, 90% sensitive – 10% resistant, 100% MCF7 resistant) for MCF7 spheroid experiments. 24 h later, spheroids were washed and fresh medium including treatment drugs was applied. Spheroids were treated for a total of 18-21 days with imaging and media change performed at every 4th and 7th day of the week. All 3D experiments were performed in DMEM complete medium except for charcoal stripped FBS medium experiments. Spheroids were treated with ribociclib (Selleck Chemicals, Cat. No: S7440), tamoxifen (Selleck Chemicals, Cat. No: S7827), raloxifene (Selleck Chemicals, Cat. No: S5781), and letrazole at specified doses described in supplementary figure 8. Imaging was performed using Cytation 5 imager (Biotek Instruments) recording signal intensity from brightfield, YFP (for Venus fluorescence) CFP 450/440 (for Cerulean fluorescence) and Texas Red (for mCherry fluorescence) channels. Raw data processing and image analysis were performed using Gen5 3.05 and 3.10 software (Biotek Instruments). Briefly, the stitching of 2 × 2 montage images and Z-projection of 6 layers using focus stacking was performed on raw images followed by spheroid area analysis. To quantify growth under these conditions, we measured fluorescence intensity and growth of spheroid area over the total time of the experiment. For cell count calculations, a standard curve was created by measuring the area of spheroids after plating at different cell numbers 24 hours after plating. A resulting equation by fitting a curve to the data was performed by GraphPad Prism 7.02 software (second order polynomial – quadratic – curve fit used). Whole spheroid area and fluorescence intensity measurements of each population were integrated into the fitted equation, and cell counts for each population were produced from fluorescence intensities relative to spheroid size. All coculture experiments were performed in triplicates.

***Western blot analysis***

Lysates of LY2 and MCF7 cells were separated by SDS-polyacrylamide gelelectrophoresis and proteins were transferred electrophoretically to a polyvinylidene difluoride membrane using Invitrogen iBlot 2 device and Invitrogen iBlot Transfer Stacks. Membranes were blocked with Tris-buffered saline with 0.05% tween 20 (TTBS) and 5% BSA for 1 hour at room temperature. After washing with TTBS, membranes were then probed with anti-HSD17β1 polyclonal antibody (Abnova, H00003292-M03A, 1:1000, overnight 4°C; R&D systems, MAB7178, 1:2000, overnight 4°C), anti- anti-HSD17β8 polyclonal antibody (Proteintech, 16752-1-AP, 1:1000 dilution, overnight 4°C), and anti-β-actin monoclonal antibody (Santa Cruz Biotechnology, sc-47778, 1:500 dilution, 1 hour room temperature) and detected using SuperSignal West Pico PLUS Chemiluminescent Substrate (Thermo Scientific) with anti-rabbit (GE Healthcare NA9341ML) or anti-mouse (GE Healthcare NXA9311ML) peroxidase-linked secondary antibody (1:6000 dilution). The molecular weight was determined using a prestained protein marker (BioRad).

***Multiplex cytokine analysis***

Media samples taken at day 21 from 3D spheroid experiments, treated with or without ribociclib, (experimental setup previously described in results and methods - ***Mono- and coculture 3D spheroid experiments***) and plated in different compositions (100% sensitive, 50% sensitive – 50% resistant, and 100% resistant) were spun down at 300g and frozen at -80 $^{o}$C. Samples were then prepared by the Analytical Pharmacology Core of City of Hope National Medical Center for multiplex cytokine analysis. Samples were analyzed for 5 cytokines (EGF, FGF2, FGF21, FGF23, and TGFα) using the "ProcartaPlex Multiplex Immunoassay Kit" (Invitrogen, Camarillo, CA) per manufacturer's protocol. In addition, TGFβ-1, -2, and -3 were measured using the "Magnetic Luminex Performance Assay Kit" (R&D Systems, Minneapolis MN) also according to manufacturer's instructions. For analysis of TGFβ, the latent proteins required activation to their immunoreactive state prior to detection. Activation was accomplished by adding 20 µl of 1N HCl to 100 µl of sample followed by incubation for 10 minutes at room temperature. Samples were then neutralized by the addition of 20 µl of 1.2N NaOH/0.5M HEPES prior to dilution and loading on to the plate. Briefly, multiplex bead solutions were vortexed for 30 s and 50 µl was added to each well and then washed twice with wash buffer. Next, 50 µl of sample (cell culture supernatant) was loaded in duplicate into Greiner flat-bottom 96 well microplates. Cytokines standards were reconstituted with unconditioned cell culture medium and serial dilutions were prepared. Plates were then incubated on a plate shaker at 500 rpm in the dark at room temperature for 2 hours. The plate was then applied to a magnetic device designed to accommodate a microplate and all wells were washed two times with 200 µl of wash buffer. Biotinylated detection antibody mix (25 µl) was added to each well and the plate was incubated on the plate shaker for another hour. After washing two times with 200 µl of wash buffer, streptavidin-phycoerythrin (50 µl) was added to each well followed by incubation on a plate shaker for 30 minutes. After two more washes, the contents of each well were resuspended in 120 µl reading buffer and shaken for 5 min. Finally, the plate was transferred to the Flexmap 3D Luminex system (Luminex corp.) for analysis, cytokine concentrations were calculated using Bio-Plex Manager 6.2 software with a five parameter curve-fitting algorithm applied for standard curve calculations for duplicate samples.

### ***Ribociclib concentration measurements with high performance liquid chromatography - mass spectrometry (HPLC/MS)***

Spheroid experiments were initiated as earlier described (***Mono- and coculture spheroid experiments*)** with some modifications. CAMA-1 sensitive and resistant cells were plated at different cell numbers (2,000; 10,000; 40,000) and in different compositions (100% sensitive, 50% sensitive – 50% resistant, and 100% resistant). After 24 hours, ribociclib treatment (400nM) was applied for 4 days. Following the 4-day treatment, media from cell samples were spun down at 300g and frozen at -80 $^{o}$C. Samples were then prepared by the Analytical Pharmacology Core of City of Hope National Medical Center for HPLC/MS. Media without cells (+/- drug) were also subjected to HPLC/MS measurements.

Acetonitrile (ACN) and methanol were of HPLC-grade and purchased from Fisher Scientific (Fair Lawn, NJ, USA). Ammonium acetate was purchased from Mallinckrodt (Kentucky, USA). Ribociclib and abemaciclib (internal standard) were provided by Selleck Chemicals. Deionized water was prepared using the Millipore Milli-Q system (Milford, MA, USA). Abemaciclib measurements were optimized to provide negative control.

Samples were prepared for analysis by mixing 30 µl of media with 10 µl 50% methanol in water in a 0.5 ml low retention micro-centrifuge tube. To this tube, 10 µl of 6 µM abemaciclib in 50% methanol and 180 µl of ice cold methanol were added. The tube was then vortex mixed for 3 minutes and centrifuged for 5 minutes at highest speed and 4°C. Following centrifugation, 20 µl of the supernatant was mixed with 180 µl of 50% methanol in 50% 10mM ammonium acetate. The final solution was then transferred to an autosampler vial and 2 µl was injected on column.

LC-MS/MS analysis was performed using a Waters Acquity UPLC system (Milford, MA, USA) interfaced with a Waters Quattro Premier XE mass spectrometer. HPLC separation was achieved using a Gemini NX C18, 100 x 2.1mm column (Phenomenex, Torrance, CA, USA) proceeded by a Phenomenex Gemini NX C18 guard column (Torrance, CA, USA). The column temperature was maintained at 40°C. Isocratic elution was performed using a mobile phase of 15% 6 mM ammonium acetate in ACN at a flow rate of 0.3 ml/min. Under optimized chromatographic conditions, the retention times were 1.2 minutes and 1.89 minutes for ribociclib and abemaciclib, respectively. The total run time was 4 minutes.

The electrospray ionization source of the mass spectrometer was operated in positive ion mode with a cone gas flow of 25 L/hr and a desolvation gas flow of 750 L/hr. Capillary voltages were

0.4 kV for both ribociclib and abemaciclib, and the cone voltages were optimized at 45 V for ribociclib and 28 V for abemaciclib, respectively. The collision voltages were set to 32 V for ribociclib and 28 V for abemaciclib. The source temperature was 125°C and the desolvation temperature was 470oC. Multiple reaction monitoring (MRM) was used for quantitation and the optimal precursor → product ion combinations were determined to be 435.34 → 321.9 m/z and 507.32 → 392.92 m/z for ribociclib and abemaciclib, respectively. MassLynx version 4.1 software was used for data acquisition and processing.

***Total exosome isolation experiment***

Spheroid experiments were initiated as earlier described (***Mono- and coculture spheroid experiments***) with some modifications. CAMA-1 sensitive and resistant cells were plated at 2,000 cells per well and in different compositions (100% sensitive and 100% resistant). These spheroids were treated with or without 400nM ribociclib and designated as donor wells. On every 3rd and 6th day of the week media was changed on the donor wells and exosomes were isolated from the old media using Total Exosome Isolation Reagent (from cell culture media, Thermo Fisher, Cat. No.: 4478359) according to the manufacturer's protocol. Briefly, cell culture media was centrifuged at 2000Xg for 30 minutes to remove cells and debris, and 200 uL supernatant was mixed with 100 uL Total Exosome Isolation Reagent and incubated on 4C overnight. Following this incubation, samples were centrifuged at 10,000Xg for 60 minutes at 4C and exosomes were resuspended in PBS. Exosomes (either from untreated or ribociclib treated, 100% sensitive or 100% resistant donor wells) were isolated from this procedure and added to the media of other 100% sensitive spheroids, designated as acceptor wells, during media change on every 4th and 7th day of the week. During this media change, 75% of media was untreated or 400nM ribociclib treated complete media with the remaining 25% contribution of isolated exosomes originating from donor wells. Imaging of each well was performed on every 4th and 7th day of the week as earlier described in methods using the Cytation 5 imaging system. Analysis was completed as previously described in methods.

***Description of FACT mechanistic model comparisons***

Using formal model comparison, we compared the accuracy of our facilitation model's predictions of spheroid growth against predictions of alternative models describing direct competition for resources (Competition alone model) or phenotypic plasticity in which cells transition from a naive to a resistant state either in response to drug induction (Plasticity DI) or via random switching (Plasticity RS). Here we outline the set of differential equation models that were used to describe alternative hypotheses regarding mechanisms that govern the growth dynamics of monocultures and cocultures of resistant and sensitive isogenic cancer cells (CAMA-1) grown under different doses of the cell cycle inhibitor ribociclib. Bayesian inference was used to fit each model to the growth trajectories of mono- and co-cultures of sensitive and resistant cells across 8 doses of ribociclib. Here we present the model comparison results showing which hypotheses about the mechanisms of cell-cell interactions were supported by the data.

***Defining alternative models describing hypothesised mechanisms governing spheroid growth dynamics***

**Competition alone**

Populations of sensitive and resistant cells compete for resources to proliferate. The abundance of cell type $i$ within a spheroid ($N_i$) depends on the balance of cell proliferation and death. Cells divide at a baseline rate ( $r_i$) which is reduced by cell cycle inhibition (ribociclib; $x$). Susceptibility to cell cycle inhibition depends on $\beta_i$. Proliferation in also reduced through competition, with each cell type having a competitive effect equal $1/K_j$. Finally cell death occurs at rate $\delta_i$. This yields the following competition model:

$$\frac{dN_i}{dt} = \frac{r_i}{(1+\beta_i x)}\left(1 - \sum_{1}^{n}\frac{N_j}{K_j}\right)N_i - \delta_i N_{i_x}$$

**Plasticity (DI): Drug induced phenotype switching**

Treatment may stimulate modified gene expression, inducing cells to transition to a more resistant state, through epigenetic reprogramming. We describe both the innately resistant and sensitive cell as transitioning from an naive state ($N_i$) to an induced resistant state ($\widetilde{N}_i$). This transition rate ($\lambda$) is proportional to drug concentration. As in the pure competition model above, proliferation depends of the ribociclib concentration and the abundance and competitiveness of naïve and induced cells of each lineage (resistant vs sensitive). Cell death occurs at differing rates in induced and un-induced cells. This yields the following drug induced phenotype switching model:

$$\frac{dN_i}{dt} = \frac{r_i}{(1+\beta_i x)}\left(1 - \sum_{j=1}^{n}\left(\frac{N_j}{K_j} + \frac{\widetilde{N}_j}{\widetilde{K}_j}\right)\right)N_i - \delta_i N_i - \lambda x N_i$$

$$\frac{d\widetilde{N}_i}{dt} = \frac{\tilde{r}_i}{(1+\tilde{\beta}_i x)}\left(1 - \sum_{j=1}^{n}\left(\frac{N_j}{K_j} + \frac{\widetilde{N}_j}{\widetilde{K}_j}\right)\right)\widetilde{N}_i - \tilde{\delta}_i \widetilde{N}_i + \lambda x N_i$$

**Plasticity (RS): Random phenotype switching**

Resistance related changes in gene expression may occur independent of treatment, for example depending on the cell cycle state that the cell happened to be in at the onset of treatment. We describe the resistant and sensitive lineages as transitioning from an naive state ($N_i$) to an induced resistant state ($\widetilde{N}_i$) at a rate that is independent of treatment ($\lambda$). Cell proliferation and death is the same as in the drug induced phenotype switching model. This yields the following random phenotype switching model:

$$\frac{dN_i}{dt} = \frac{r_i}{(1+\beta_i x)}\left(1 - \sum_{j=1}^{n}\left(\frac{N_j}{K_j} + \frac{\widetilde{N}_j}{\widetilde{K}_j}\right)\right)N_i - \delta_i N_i - \lambda N_i$$

$$\frac{d\widetilde{N}_i}{dt} = \frac{\tilde{r}_i}{(1+\tilde{\beta}_i x)}\left(1 - \sum_{j=1}^{n}\left(\frac{N_j}{K_j} + \frac{\widetilde{N}_j}{\widetilde{K}_j}\right)\right)\widetilde{N}_i - \tilde{\delta}_i \widetilde{N}_i + \lambda N_i$$

**Facilitation Symmetric (1D): Allee effect model of facilitation**

We first describe facilitation between resistant and sensitive cells in which cells of both lineages contribute equally to the facilitation effect on a per cell basis (symmetric facilitation effects). We describe the production of a facilitation factor that is produced at rate $\epsilon$ in both cell types. The beneficial effect of the facilitation factor saturates at high concentrations, with the asymptotic benefit at high densities equalling $1/\ c$. This facilitation effect increases proliferation, whilst the functional form of competition, the drug impact and cell death are the same as in the competition model. This yields the following Allee effect model of symmetric facilitation:

$$\frac{dN_i}{dt} = \frac{r_i}{(1+\ \beta_i x)}\left(1 - \sum_{j=1}^{n}\frac{N_j}{K_j}\right)\left(1 + \frac{\epsilon \sum_{j=1}^{n} N_j}{1 + c\ \sum_{j=1}^{n} N_j}\right)N_i -\ \delta_i N_{i_x}$$

**Facilitation (1D): Asymmetric contribution to facilitation**

The production and decay of a facilitation factor ($E_E$) can be modelled explicitly to account for the differences in production by sensitive and resistant cells. Cells of cell type $i$ produce the a facilitation factor at rate $\gamma_j$ and it decays at rate $\delta_E$. The concentration of the facilitation factor determines the proliferation promotion benefit. As with the symmetric facilitation model, this benefit saturates at high concentrations, with an asymptotic benefit of $1/\,c$. The functional form of competition, the drug impact and cell death are again the same as in the basic competition model. With $E_E$ in quasi steady state and symmetric contributions of cell types to facilitation, this reduces to the Allee effect model of facilitation.

$$\frac{dN_i}{dt} = \frac{r_i}{(1+\ \beta_i x)}\left(1 - \sum_{j=1}^{n}\frac{N_j}{K_j}\right)\left(1 + \frac{E_E}{1 + c\ E_E}\right)N_i -\ \delta_i N_{i_x}$$

$$\frac{dE_E}{dt} = \sum_{j=1}^{n}\gamma_j N_j -\ \delta_E E_E$$

**Facilitation (2D): Asymmetric contribution to facilitation & cell quiescence**

To describe the mechanism of action of ribociclib treatment more mechanistically, we use the stage-structured modelling approach to create a minimal population level model to predict the effects of cell cycle inhibition treatment (1).

With this approach, cells transition from a proliferative ($P$), to a quiescent ($Z$) state. Proliferative cells enter the G1/S phase cell cycle checkpoint at a baseline rate ($r$), which is reduced by resource competition ($\alpha(P,Z,X)$) and increased by estradiol availability ($E_E$). Resource competition between cells is described by $\alpha(P,Z) = 1 -\ \sum_{j=1}^{n}\frac{P_j +\ Z_j}{K_j}$, where competitive ability of each cell type ($j$) is determined by the carrying capacity parameter $K_j$. As in the 1D facilitation model, the extracellular concentration of the facilitation factor ($E_E$) depends on its differing net production by sensitive and resistant cells ($\gamma_j$) and its decay ($\delta_E$). Additionally, we describe the intracellular concentration of facilitation factors, which depends on the balance of production by the cell and influx into the cell against diffusion out of the cell and binding. This leads to the intracellular steady state concentration of cell type $i$ of: $E_i =\ \frac{\rho_i + \eta E_E}{\eta +\ \mu_i}$.

Cell entry into the G1/S phase increases with the intracellular level of estradiol ($E_i$) and increased receptor binding ($\mu_i$), saturating at high concentrations when uptake and binding becomes rate limited ($c$). We describe this as: $G_i = r_i(1 + \frac{\mu_i E_i}{1 + c\mu_i E_i})\alpha(P,Z)$. Cells in the G1/S phase undertake a decision to divide or enter a quiescent state, based on the balance of key regulatory cell cycle promoters and inhibitors. Cell cycle inhibition using ribociclib ($x$) inactivates key promoters of the G1/S checkpoint transition, blocking cell cycle progression and increasing the probability of quiescence above the baseline ($\lambda_i$) according to $q_i(x) = \frac{x}{k_i+x}$, where the half-saturation constant ($k_i$) can differ between resistant and sensitive cells to quantify drug susceptibility. Following quiescence, cell death occurs at rate $\varphi_i$.

Together these components capturing competition, facilitation, cell cycle progression and arrest yield the population model:

$$G_i = r_i\left(1 - \sum_{j=1}^{n} \frac{P_j + Z_j}{K_j}\right)\left(1 + \frac{\mu_i E_i}{1 + c_i \mu_i E_i}\right)$$
$$q_i(x) = \frac{x}{k_i + x}$$

$$\frac{dP_i}{dt} = \left( G_i\big(1 - q_i(x)\big) - G_i q_i(x) - \lambda_i\right)P_i$$
$$\frac{dZ_i}{dt} = (G_i q_i(x) + \lambda_i)P_i - \varphi_i Z_i$$

$$\frac{dE_E}{dt} = \left(\sum \gamma_i (P_i + Z_i)\right) - \delta_E E_E.$$

**Facilitation (3D): Asymmetric contribution to facilitation & cell quiescence and senescence**

Finally, we extend the stage structured model to describe the transition from proliferative ($P$), to quiescent ($Z$) and senescent ($X$) cell states.

Proliferative cells enter the G1/S phase cell cycle checkpoint at a baseline rate ($r$), which is reduced by resource competition ($\alpha(P, Z, X)$) and increased by estradiol availability ($E_E$). Resource competition between all three cell states extends to: $\alpha(P, Z, X) = 1 - \sum_{j=1}^{n} \frac{P_j + Z_j + X_j}{K_j}$, where competitive ability of each cell type ($j$) is determined by the carrying capacity parameter $K_j$. As above, cell entry into the G1/S phase increases with the intracellular level of estradiol ($E_i$) and increased binding ($\mu_i$), saturating at high concentrations when uptake and binding becomes rate limited ($c$), which we describe as: $G_i = r_i(1 + \frac{\mu_i E_i}{1 + c\mu_i E_i})\alpha(P, Z, X)$. Cells in the G1/S phase either divide or enter a quiescent state, following the same mechanism as described in the 2D facilitation model. However, following quiescence, cells transition into the final senescent state at rate $\varphi_i$ before cell death occurs at rate $\delta_i$. Although structurally and mechanistically similar, this model produces additional delays in the drug effects on population abundances. This full model and its derivation is described in more detail in the methods section but follows:

$$G_i = r_i\left(1 - \sum_{j=1}^{n} \frac{P_j + Z_j + X_j}{K_j}\right)\left(1 + \frac{\mu_i E_i}{1 + c_i \mu_i E_i}\right)$$
$$q_i(x) = \frac{x}{k_i + x}$$

$$\frac{dP_i}{dt} = ( G_i\left(1 - q_i(x)\right) - G_i q_i(x) - \lambda_i)P_i$$

$$\frac{dZ_i}{dt} = (G_i q_i(x) + \lambda_i)P_i - \varphi_i Z_i$$

$$\frac{dX_i}{dt} = \varphi_i Z_i - \delta_i X_i$$

$$\frac{dE_E}{dt} = \left(\sum \gamma_i(P_i + Z_i)\right) - \delta_E E_E.$$

***Results of models comparison showing which hypotheses were supported by the data***

Formal probabilistic model comparison of the non-nested alternative hypotheses was performed using Watanabe–Akaike information criterion (WAIC). This sums the average fit (log likelihood) of posterior samples over all data points and penalizes the complexity of the model hypothesis based on the pointwise variance in the fit across posterior uncertainty (55). The resulting WAIC score measures the goodness of fit of the model to the data whilst penalizing for model complexity to avoid overfitting. Models incorporating facilitation greatly outperformed (lower penalized prediction error) models of competition alone or phenotypic plasticity. Facilitation models assuming equivalent facilitation by resistant and sensitive cells (Facilitation symmetric 1-State) were greatly outperformed by facilitation models in which resistant cells contribute disproportionately to the production of estradiol (Facilitation 1/2/3-State). The addition of quiescent and senescent cell states further improved model accuracy compared to simpler models in which ribociclib simply reduces cell division rates (Facilitation 2/3-State versus Facilitation 1-State). These additional states allow the prediction of the delayed impacts of therapy at higher doses (reducing penalized prediction error) and also the estimation of the rate of division and quiescence with and without therapy. Describing cells as transitioning through proliferative, quiescent and senescent states during the cells life (Facilitation 3-State; model schematic in **Fig. S10**) yielded a superior description of the data after penalizing for model complexity. This analysis guarded against model overfitting and showed the explanatory power of the estradiol mediated facilitation hypothesis and the model of this process that we present in the main text.
